# Identifying Y-chromosome haplogroups in arbitrarily large samples of sequenced or genotyped men

**DOI:** 10.1101/088716

**Authors:** G. David Poznik

## Abstract

We have developed an algorithm to rapidly and accurately identify the Y-chromosome haplogroup of each male in a sample of one to millions. The algorithm, implemented in the yHaplo^*^ software package (yHaplo), does not rely on any particular genotyping modality or platform. Full sequences yield the most granular haplogroup classifications, but genotyping arrays can yield reliable calls, provided a reasonable number of phylogenetically informative variants has been assayed. The algorithm is robust to missing data, genotype errors, mutation recurrence, and other complications. We have tested the software on full sequences from phase 3 of the 1000 Genomes Project and on subsets thereof constructed by downsampling to SNPs present on each of four genotyping arrays. We have also run the software on array data from more than 600,000 males.

## Introduction

As the Y chromosome bears the longest stretch of non-recombining DNA in the human genome, it contains sufficient information to reconstruct a detailed phylogenetic tree relating the lineages of every man to the most recent common male-line ancestor of all men^1^. The clades of this tree are called “haplogroups,” a portmanteau of “haplotype” and “group.” Haplogroups are highly correlated with geography and population, and their global distributions have lent much insight into historical migrations^2^.

We have developed yHaplo, a tool that enables researchers to rapidly and accurately identify the Y-chromosome haplogroup of each male in a genetic sample of arbitrary size. This tool will be particularly useful for large-scale Y-chromosome studies, as it will allow the analyst to partition the sample prior to tree inference with no loss of information. But it will also be useful for more general population-genetic or medical-genetic studies in which Y-chromosome data is often generated incidentally. For these studies yHaplo will help the analyst to quickly and easily assess which paternal lineages are present in their study sample. This knowledge could, for example, indicate the presence of undocumented admixture that primary analyses should take into account^3^.

At the time of writing, yHaplo can assign 1,757 unique haplogroup labels, but it can easily be updated to reflect future refinements of the tree, as it reads from two simple text files all information about the structure of the tree and the markers that define it.

## Methods

yHaplo builds its representation of the Y-chromosome phylogeny in two steps. First, it imports a text file that explicitly codifies the primary structure of the tree—the major haplogroups and their relationships to one another (Figure 1). It then reads in a set of phylogenetically informative single nucleotide polymorphisms (SNPs) curated by the International Society of Genetic Genealogy (ISOGG)^4^ and associates each with a specific branch, growing the tree as necessary according to the nomenclature convention delineated by the Y-Chromosome Consortium^5^. For example, to store the SNP named M343, which is a haplogroup-R1b defining C→A mutation at coordinate 2,887,824, the software creates two children of haplogroup R, naming them R1 and R2, and then creates two children of R1, naming them R1a and R1b. It then associates M343 with the newly formed R1b branch.

**Figure 1.**
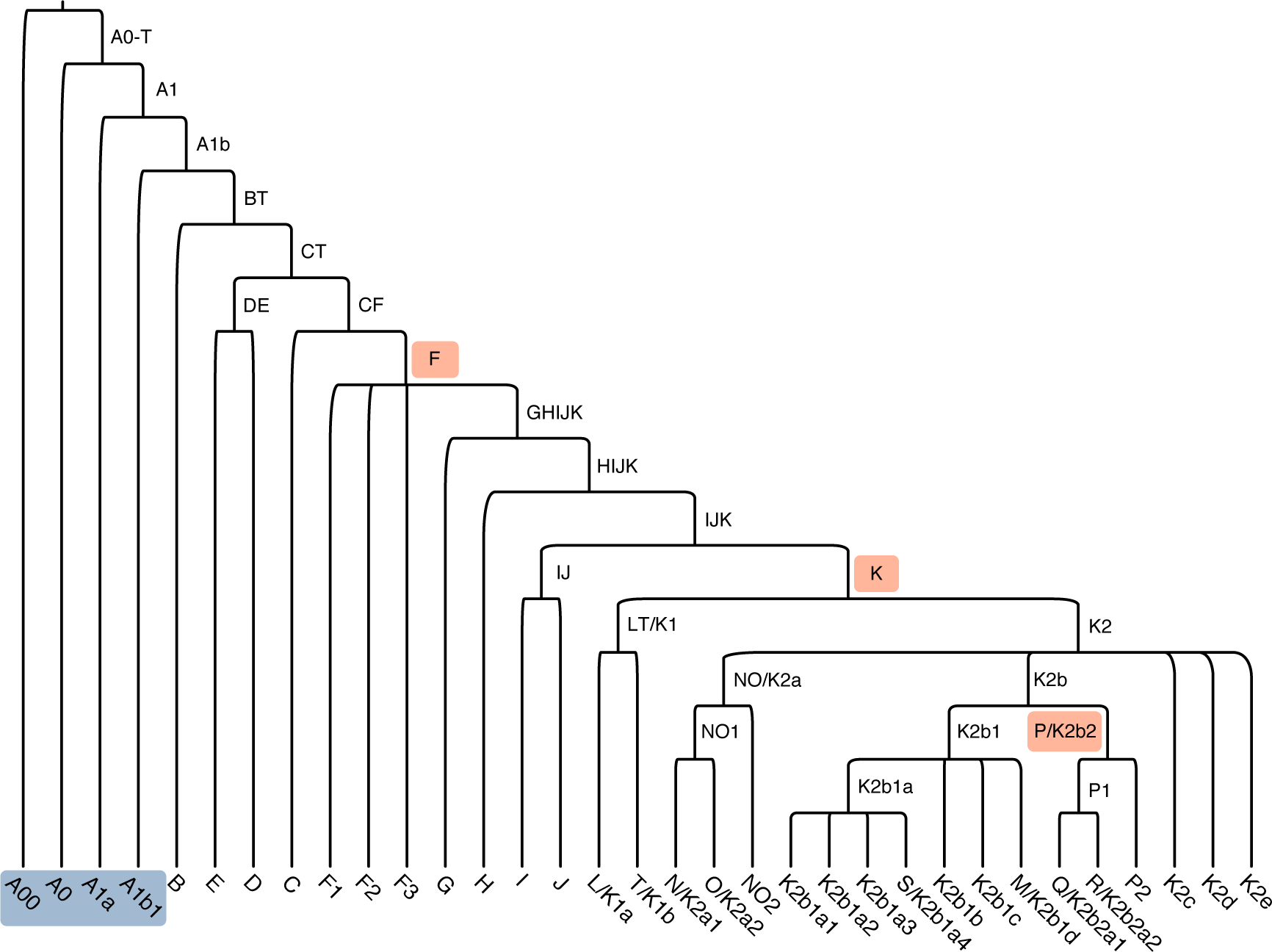
Primary structure of the Y-chromosome tree. Nineteen letters label monophyletic clades, but three of these denote internal branches ancestral to other haplogroups (orange): F is an ancestor of G, H, I, J, and K; K is the common ancestor of L, T, N, O, S, M, and P; and P is an ancestor of Q and R. A twentieth letter, “A,” marks a paraphyletic group of the four most highly diverged clades: A00, A0, A1a, and A1b1 (blue). Multi-letter labels represent joins. For example, DE is the parent of D and E. Finally, A1b is the parent of A1b1 and BT, the common ancestor of all non-A haplogroups.

To call an individual’s haplogroup, we search for a path of branches through the Y-chromosome tree from its root to the most terminal branch possible. With artificially ideal data, we could conduct a simple breadth-first search, as we would expect to observe exclusively derived-state genotypes for the SNPs associated with branches along the correct path and exclusively ancestral-state genotypes elsewhere. However, we must contend with missing data, genotype errors, mutation recurrence (homoplasy), and undetected errors in the database of phylogenetically informative SNPs. Therefore, we have designed a modified breadth-first search that is robust to these complications.

To identify the evolutionary path leading to an individual’s lineage, we first initialize a queue of search paths, each of which tracks the derived alleles observed thus far and specifies the next branch to consider. At first, the queue consists of two paths: one for each of the two branches that arose with the tree’s most ancient branching event. We then iterate through the queue until it is empty.

Upon dequeuing an active path, we assess the individual’s genotypes for all SNPs associated with the next branch, noting the numbers of alleles observed in the ancestral and derived states, and ignoring SNPs for which no data is available for the individual. We then either put a stop to the search path or continue it by forking—replicating it for each child of the current branch and enqueuing each replicate to continue exploring. For efficiency, whenever the current path is clearly the correct one, as indicated by a large number of derived-allele genotypes, we wipe the queue clean prior to forking.

The search path stops on its own if the branch under examination is terminal (i.e., if it leads to a leaf node). Otherwise, to decide whether to stop or to continue, we consider the number of ancestral and derived alleles (Table 1). If we observe no ancestral alleles, there is no evidence suggesting we should stop the search, so we fork, whether or not we observe any derived alleles. Similarly, we continue pursuing a path when we observe just one ancestral allele, as it could be due to a genotyping error or to a reversion mutation. Conversely, we deem three or more ancestral alleles compelling evidence to stop a line of inquiry, regardless of the number of derived alleles we observe on the branch. Finally, if we observe exactly two ancestral alleles, we hinge the decision on the number of derived alleles. If we observe none, the individual carries the ancestral allele at the only two sites associated with this branch, and we regard this as sufficient evidence to stop. However, when we observe one or more derived alleles in addition to the two ancestral alleles, we continue the search in case both ancestral alleles were errors or reversions. yHaplo users can easily adjust any of the parameters described above, but we have found that they work quite well.

**Table 1.** Search stopping conditions. **# Anc** and **# Der**, number of ancestral and derived alleles observed on a branch.

| # Anc | # Der | Stop | Reason |
| --- | --- | --- | --- |
| 0, 1 |  | No | Insufficient evidence to stop, or none at all |
| 2 | 1+ | No | Given evidence to continue, do so for robustness |
|  | 0 | Yes | Reasonable evidence to stop and no evidence to continue |
| 3+ |  | Yes | Strong evidence to stop |

We defined the stopping conditions to be robust to a particularly challenging case, when just a single SNP is genotyped on a branch, and the observed genotype corresponds to the ancestral allele due to an error or reversion mutation. On its own, this permissiveness could be exploited, particularly when a genotyping platform sparsely covers regions of the tree. Therefore, we track whether or not a path has pushed through a branch with one ancestral and zero derived SNPs. If it has, and subsequent steps in the path find no evidence for downstream derived alleles, we revert it in a post-processing step.

When the queue is empty, we compare the stopped paths and select the one with the most SNPs observed in the derived state. This best path indicates the individual’s haplogroup.

## Results

We tested yHaplo on the Y-chromosome data from phase 3 of the 1000 Genomes Project^3,6^ (1000G). These data include full-sequencing-derived genotypes at ~60,000 single nucleotide variants for 1,244 males from 26 globally diverse populations. We constructed a gold-standard set of haplogroup calls using a semi-automated method^3,7^ and an up-to-date version of the of the ISOGG database. As this method requires a manual curation step, it is somewhat onerous to apply to samples on the order of 1,000 or larger, but it yields extremely reliable calls, and the haplogroup assignment emitted by yHaplo is identical to the gold-standard call for each 1000G male.

For each of the four versions of the 23andMe genotyping array, we downsampled the 1000G data to the set of SNPs present on the platform and re-ran the software. The resulting haplogroup calls were of course less granular than those based on the full sequencing data, but each was accurate.

We have run yHaplo on more than 600,000 male customers in the 23andMe database and manually inspected the data to confirm haplogroup calls for several hundred individuals, with particular emphasis on those bearing lineages with relatively few diagnostic markers assayed.

## Discussion

We have described an algorithm, and a Python implementation thereof, that leverages the tree-structure of Y-chromosome diversity to rapidly deduce accurate haplogroup assignments for large samples of sequenced or otherwise genotyped men. This method is robust to genotype errors, recurrent and reversion mutations, and undetected errors in the input database of phylogenetically informative SNPs. It uses current nomenclature and knowledge of the Y-chromosome phylogeny. As it reads from two simple text files all information about the structure of the tree and the SNPs that define it, calls can easily be updated to reflect future refinements.

When using its preferred sample-major genotype data format, yHaplo processes the 1,244 male sequences from phase 3 of the 1000 Genomes Project in 20 seconds, and in under 1.5 minutes, it can run directly on the SNP-major VCF file hosted by the Project. Finally, the software includes more than 20 command-line options to construct a variety of informative auxiliary output files that summarize the haplogroup calling process, features of the tree, and the degree to which the tree is covered by any particular genotyping platform. We describe many of these options in the accompanying software manual.

## Software

yHaplo is available at the 23andMe code repository for non-commercial use pursuant to the terms of the non-exclusive license agreement included with the software distribution:

https://github.com/23andMe/yhaplo

## Acknowledgements

We thank Kasia Bryc and Joanna Mountain for proposing the project and for helpful discussions.

## Footnotes

* yHaplo is a trademark of 23andMe, Inc.

## References

1. Underhill, P. A. et al. Y chromosome sequence variation and the history of human populations. Nat. Genet. 26, 358–61 (2000).

2. Chiaroni, J., Underhill, P. A. & Cavalli-Sforza, L. L. Y chromosome diversity, human expansion, drift, and cultural evolution. Proc. Natl. Acad. Sci. 106, 20174–20179 (2009).

3. Poznik, G. D. et al. Punctuated bursts in human male demography inferred from 1,244 worldwide Y-chromosome sequences. Nat. Genet. 48, 593–9 (2016).

4. ISOGG. International Society of Genetic Genealogy. (2016). at <http://www.isogg.org/>

5. The Y Chromosome Consortium. A nomenclature system for the tree of human Y-Chromosomal binary haplogroups. Genome Res. 12, 339–348 (2002).

6. The 1000 Genomes Project Consortium. A global reference for human genetic variation. Nature 526, 68–74 (2015).

7. Poznik, G. D. et al. Sequencing Y chromosomes resolves discrepancy in time to common ancestor of males versus females. Science 341, 562–565 (2013).

